# LabGrymace: Automated Analysis of Mouse Grimace for Quantitative Assessment of Pain Dynamics

**DOI:** 10.64898/2026.07.31.742042

**Authors:** Wenjin Dong, Kayla M. Moehn, Elizabeth A. Ronan, Yujia Hu, Joshua J. Emrick, Bing Ye

## Abstract

Accurate assessment of pain in animal models is essential for understanding pain mechanisms, developing analgesics, and ensuring animal welfare. The Mouse Grimace Scale (MGS) provides a sensitive, non-invasive measure of spontaneous pain by quantifying pain-related facial expressions, but its utility is limited by labor-intensive manual scoring, observer variability, and reliance on static images that fail to capture the temporal dynamics of facial behavior. Existing automated approaches improve throughput but typically rely on highly standardized imaging conditions, selected viewing angles, and static facial appearance, while providing limited temporal resolution and little insight into the relative contributions of individual facial action units. Here, we introduce **LabGrymace**, an open-source, artificial intelligence-powered framework for automated, frame-by-frame analysis of pain-related facial dynamics in freely moving mice. Built on the LabGym behavioral analysis platform, LabGrymace uses deep-learning-based facial feature detection and tracking to quantify ear, eye, and nose movements continuously from video recordings. To generate a quantitative pain metric and facilitate reproducibility, we calibrated facial dynamics against graded chemogenetic activation of nociceptors and identified the kinematic features most strongly associated with pain intensity. These features were integrated into a weighted composite pain score that reflects the differential contributions of individual facial action units. LabGrymace accurately classified pain-related facial actions and generated continuous pain scores without manual frame selection or restrictive recording conditions. The resulting pain scale exhibited dose-dependent responses generalized across distinct pain modalities, including visceral pain induced by MgSO₄ and somatic pain induced by capsaicin. By combining automated facial-feature analysis with quantitative temporal modeling, LabGrymace provides an objective, scalable, interpretable, and flexible tool for assessing spontaneous pain in laboratory mice.

## INTRODUCTION

Accurately assessing pain in model organisms is critical for identifying novel pain mechanisms, discovery of analgesics, and ensuring animal welfare. Historically, researchers have relied on withdrawal responses to mechanical, thermal, or electrical stimulation to quantify nociceptive responses. While these methods are valuable for measuring sensitivity to a given stimulus, they may fail to capture the dynamics of affective and perceptual dimensions of pain.

To capture pain states, the development of the Mouse Grimace Scale (MGS) revolutionized preclinical pain research, demonstrating that mice exhibit highly reproducible facial expressions after chemical, inflammatory, or surgical insult (Langford et al. 2010). By quantifying changes across distinct facial action units (e.g., eye orbital tightening, nose bulging, cheek bulging, ear position, and whisker position), the MGS provides a sensitive, non-invasive readout of a mouse’s internal pain state. However, the traditional application of the MGS relies heavily on manual scoring by trained observers. This approach is not only labor-intensive and low-throughput but also suffers from subjective variability among experimenters and laboratories. Furthermore, manual assessment usually requires acquiring static snapshots or short video clips, which may fail to capture the dynamic, temporal structure of spontaneous facial movements.

Machine learning approaches are attractive for automation of animal behavior analysis because they may reduce user workload and improve reproducibility (Kabra et al. 2013, Wiltschko et al. 2015, Mathis et al. 2018, Graving et al. 2019, Hsu et al. 2021, Hu et al. 2023, Goodwin et al. 2024, Goss et al. 2024). Several tools have been developed to automate mouse facial grimace analysis, moving the field beyond manual MGS scoring and toward more scalable facial pain assessment. Tuttle et al. (2018) showed that convolutional neural networks (CNNs) could reproducibly distinguish “pain” from “no-pain” in albino mice (Tuttle et al. 2018). Later systems, such as DeepMGS (Chiang et al. 2022), moved beyond binary classification to predict individual facial action unit scores. PainFace further improved scalability and throughput by evaluating sampled video frames individually and summarizing scores across time windows (McCoy et al. 2024). GrimACE extended automated pain assessment to cage-side recordings of freely moving mice (Sturman et al. 2026). Together, these studies established the value of automation for throughput, objectivity, and reproducibility.

Despite these advances, existing automated pain-scoring approaches share at least five key areas for improvement:

(1) Rigid recording conditions. Most methods require highly standardized recording conditions, head-fixation, or specific viewing angles optimized for particular datasets or pain models. For example, PainFace requires specially illuminated imaging chamber and frontal facial views in which the eyes, ears, nose, and whiskers are all clearly visible (McCoy et al. 2024). In freely moving animals, this often necessitates labor-intensive frame selection or video trimming to obtain a sufficient number of informative facial views.

(2) Limited temporal resolution. As with conventional MGS scoring, existing methods provide only limited access to the continuous dynamics of pain-related facial behavior. Although PainFace tracks changes in pain over time, it analyzes only one frame per second rather than every frame (McCoy et al. 2024), limiting its ability to capture rapid facial dynamics. Combined with its requirement for complete facial visibility, stable, continuous acquisition of informative facial features during unrestricted movement remains challenging.

(3) Restricted quantitative resolution. Most existing tools report coarse ordinal or binary outputs, offering limited resolution for quantifying pain levels.

(4) Subjective pain calibration. Most existing models are trained to reproduce subjective grimace ratings assigned by experimenters (Chiang et al. 2022, Arnold et al. 2023, McCoy et al. 2024) or are anchored to binary states (Tuttle et al. 2018, Kobayashi et al. 2025). Consequently, their outputs inherit the subjectivity and high inter-rater variability of manual scoring, limiting reproducibility across experimenters and laboratories.

(5) Equal weighting of facial action units. Most approaches assume that all facial action units contribute equally to pain assessment (Langford et al. 2010, Sotocinal et al. 2011, Matsumiya et al. 2012, Hager et al. 2017, Chiang et al. 2022, Arnold et al. 2023, McCoy et al. 2024, Sturman et al. 2026), despite evidence that different facial features vary in their sensitivity to pain.

Consequently, the preclinical pain field still lacks an open, flexible, and interpretable framework for continuous, quantitative analysis of pain-related facial dynamics in freely moving mice. A system capable of analyzing facial movements frame by frame, quantifying pain with high temporal resolution, and objectively integrating the contributions of individual facial action units would overcome these limitations and facilitate studies of pain mechanisms, neural circuit function, and therapeutic interventions.

To bridge this gap, we applied LabGym to detect and quantify the facial dynamics in preclinical mouse models of pain. LabGym is an open-source, artificial intelligence (AI)-powered behavioral analysis platform designed for user-defined behavioral categorization and multi-parameter quantification across diverse animal species (Hu et al. 2023, Goss et al. 2024). Unlike traditional tracking software confined to rigid geometric points or coordinate tracking, LabGym utilizes deep learning to perform holistic spatiotemporal assessments. It can be trained to recognize custom behavior patterns from raw video data and, critically, computes both unique temporal descriptors such as the count, duration, latency, and intensity of a given behavior in addition to kinematic measurements including speed, velocity, acceleration, distance, magnitude area, magnitude length, vigor, intensity area and intensity length. Together, LabGym provides a granular, reproducible footprint of animal actions. This foundation has already been shown to be relevant to pain research, where LabGym has been used to quantify whole-animal spontaneous and naturalistic pain-related behaviors in an automated and holistic manner (Ronan et al. 2025, Muwanga et al. 2026).

In this study, we introduce **LabGrymace**, a specialized application of the LabGym framework tailored specifically to analyze the facial expressions of freely moving mice. We trained customizable neural networks within LabGym on facial recordings. We further developed a quantitative pain score metric for precise pain evaluation. LabGrymace can continuously detect and score pain-induced facial changes in freely moving animals, frame-by-frame without the need for manual image selection. We demonstrate that LabGrymace reliably distinguishes dose-dependent pain intensity across different nociceptive models. By capturing the full temporal dynamics of facial expression via LabGym’s framework of robust kinematic metrics, LabGrymace provides an objective, scalable, flexible, and highly interpretable open-source tool for the automated assessment of pain in laboratory mice.

## RESULTS

### The pipeline of LabGrymace

To establish LabGrymace for the analysis of facial expressions, we used the Detector and Categorizer modules within the LabGym graphical user interface (GUI). The Detector, introduced in LabGym2 (Goss et al. 2024), generated segmentation masks for user-defined facial features, enabling robust tracking even in the presence of background noise or changes in posture. Videos were first trimmed and cropped to a size that encompasses the entire body of the animal (Figure 1A). Subset of frames were then extracted from the preprocessed videos for manual annotations of facial features and used to train the Detector (Figure 1B). We employed object-annotation tools Roboflow (Dwyer et al. 2026) and EZannot (Hu 2026) to perform AI-assisted annotation of facial features in training images. Roboflow was used for labelling while EZannot was used for image augmentations. These annotated images were then used to train a Detector model to identify the ear, eye, and nose of a mouse (Figure 1C). Once trained, the Detector was applied within LabGym to extract and track facial regions from previously unseen video recordings.

**Fig. 1.**
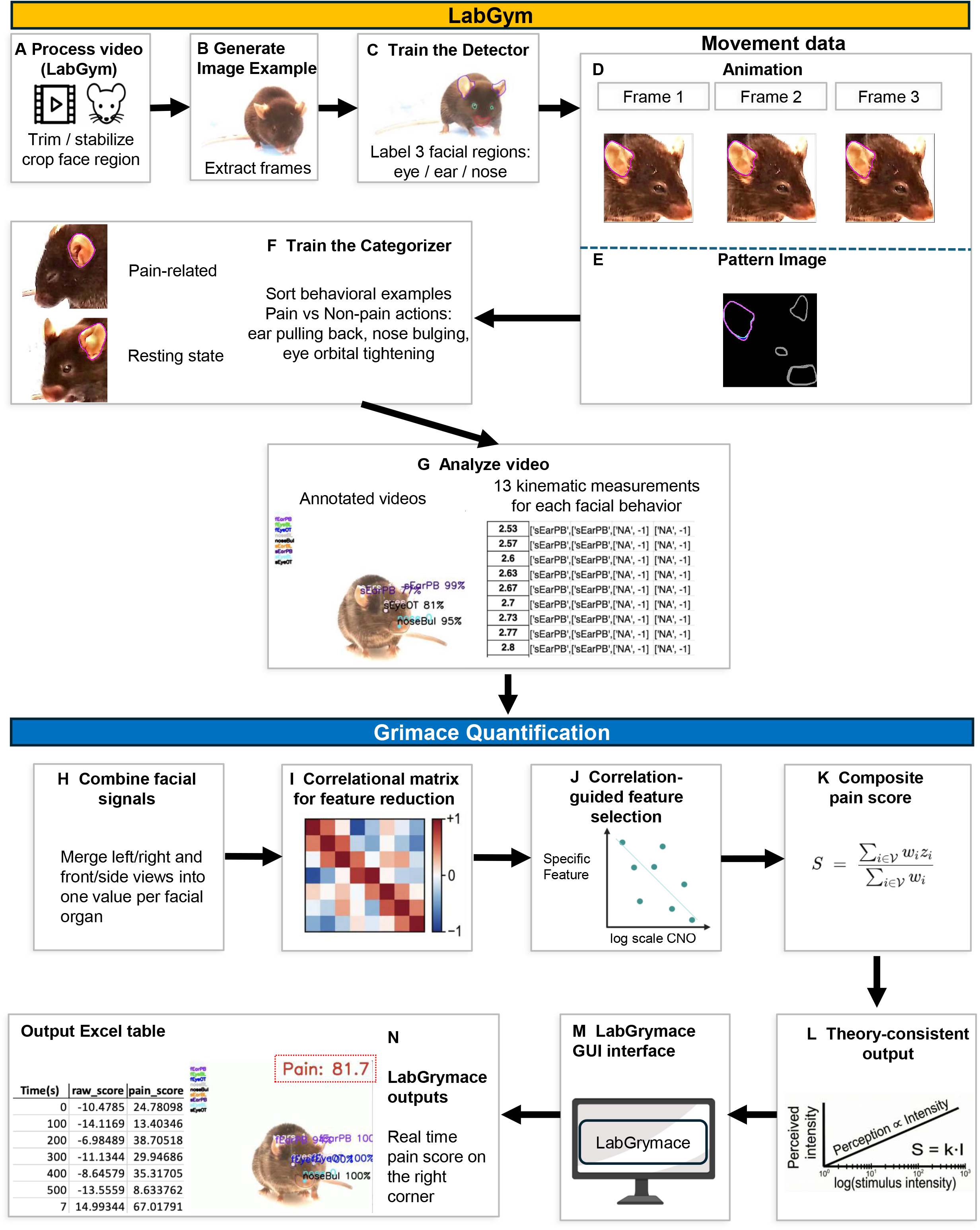
LabGrymace pain scoring pipeline. **(A)** Videos are trimmed, stabilized, and cropped to isolate the facial region of each mouse. **(B)** Image frames are extracted from behavioral videos for subsequent labelling and detector training. **(C)** A detector is trained on labeled facial features corresponding to the eye, ear, and nose. **(D)** LabGym generates an animation overlaying the facial regions’ contour across consecutive frames, capturing the temporal dynamics of facial movement. **(E)** LabGym generates a pattern image encoding the animal’s facial organs’ contour and spatial configuration, representing static morphological features. The animation (C) and pattern image (D) together constitute a behavioral example used to train the Categorizer. **(F)** A Categorizer is trained to assign facial regions’ instances to six classes spanning pain-related and resting-state facial behaviors. **(G)** LabGym analyzes the video and exports 13 kinematic measurements for each facial behavior. **(H)** Take “analyzed videos” as input and generate summary files in preparation for the subsequent steps. **(I)** Correlation matrices are used to identify redundant features within each facial region. **(J)** Selection based on correlation retains the most informative feature set for each facial region. **(K)** The composite pain score S is calculated as a weighted average of z-score-normalized facial features. **(L)** The resulting score is evaluated against Weber-Fechner-like scaling and used for downstream group comparisons. **(M)** LabGrymace graphical user interface (GUI). Users can import LabGym behavioral analysis outputs into the interface, which applies the pain scoring algorithm to generate a quantitative pain report exported as an Excel file, including per-frame pain scores and their changes over time. The GUI also supports overlaying the computed pain scores directly onto the original video for visualization. **(N)** Representative output of the LabGrymace overlay. The computed pain score, derived from the weighted pain scoring algorithm, is displayed in real time at the upper right corner of the video frame. The predicted probability that each detected facial action unit reflects a pain-related state is also overlaid on the frame (e.g., fEarPB 98%, fEyeOT 98%, noseBul 100%).

Once facial regions were identified using the Detector, LabGym generated movement data for each facial region across consecutive, user-defined time windows, which capture dynamic changes in facial morphology and movement. The movement data consisted of an animation over the user-defined time window (Figure 1D) and a paired 2-D “pattern image” summarizing position changes over time (Figure 1E) (Hu et al. 2023).

The movement data from training datasets served as the behavioral examples used to train the LabGym Categorizer. We sorted these behavioral examples into user-defined categories, including orbital tightening, ear position changes, and nose bulging. Because LabGym employs supervised learning, this annotation step provided the labeled examples required for the model to learn associations between facial movements and pain.

The labeled behavior examples were then used to train the Categorizer, which employs deep neural networks to analyze the temporal dynamics of facial movements (Figure 1F). Subsequently, the Analysis module used the trained Categorizer to extract 13 quantitative measurements. Nine of the 13 measurements were relevant to kinematic features for LabGrymace, including the acceleration, intensity area, intensity length, magnitude area, magnitude length, speed, velocity, vigor area, and vigor length (see definitions in Materials and Methods), as they reflect the temporal dynamics of an animal’s facial changes (Figure 1G).

The Categorizer classified frontal and side views as separate classes, which were merged together with the left-side and right-side signals into a single value per facial region (see Materials and Methods) (Figure 1H). From the combined quantitative measurements of facial-region movements generated by LabGym, we identified the most robust kinematic descriptors shared across facial regions, as well as their relative contributions (Figure 1I&J), and integrated these into a composite pain score (Figure 1K). During this process, we removed correlated and redundant measurements through feature-correlation analysis. We retained the remaining high-variance measurements to maximize sensitivity to pain-level differences (Figure 1L). Finally, feature weights were determined from the slope of dose-response relationships, enabling the generation of a fully quantitative pain score that scales with stimulus intensity (Figure 1L).

We designed the LabGrymace graphic user interface (GUI) (Figure 1M) to allow generation of (1) frame-by-frame pain scores, (2) temporal plots of facial-expression dynamics, (3) annotated videos in which tracked facial features and corresponding pain scores are overlaid onto the original recordings (Figure 1N), and (4) a quantitative pain report exported as an Excel file, including per-frame pain scores and their changes over time (Figure 1N).

### LabGym accurately analyzes pain-related facial action units

Unlike static image-based or frame-selected approaches, continuous video-based analysis of freely moving mice requires facial actions that remain resolvable across rapid motion and large changes in head angle. We focused on three of the pain-related facial action units from the MGS—ear position, orbital tightening, and nose bulging (Langford et al. 2010)—which were the most consistently discernible and classifiable facial features in our recordings and correspond directly to canonical MGS-associated responses. By contrast, although whisker and cheek movements are also included in the MGS (Langford et al. 2010) and may contain pain-relevant information, these features were not reliably distinguishable in black-coated mice, even by human observers, under low-contrast conditions and varying viewing angles. Including these features into the video-based pipeline would therefore be more likely to increase computational burden and classification noise than to improve pain quantification.

In the MGS, *orbital tightening* is defined as narrowing of the orbital area with eyelid closure, *nose bulging* as a rounded extension of skin visible on the bridge of the nose, and *ear position* as ears pulled apart and backward from baseline, often accompanied by vertical ridges formed as the ear tips retract (Langford et al. 2010).

To classify facial action units, we trained a LabGym Categorizer, *LabGrymace_categorizer*, to distinguish 10 facial-state classes spanning frontal and side views: the ear in the baseline position (fEarBL, sEarBL), the ear pulled back (fEarPB, sEarPB), the eye in the baseline position (fEyeBL, sEyeBL), orbital tightening (fEyeOT, sEyeOT), the nose in the baseline position (noseBL), and the nose bulging (noseBul). We manually sorted labeled examples into these 10 classes and divided them into training (8,619 examples; 80%) and validation (2,155 examples; 20%) sets. To evaluate the Categorizer’s performance under the same data distribution used for training, the validation dataset was augmented using the same pipeline as the training dataset. On this augmented validation set, the Categorizer achieved an overall accuracy of 0.97 and per-class F1 scores ranging from 0.87 to 0.99 (Figure 2G,H), indicating that the selected facial action units were classified robustly across front and side views.

**Fig. 2.**
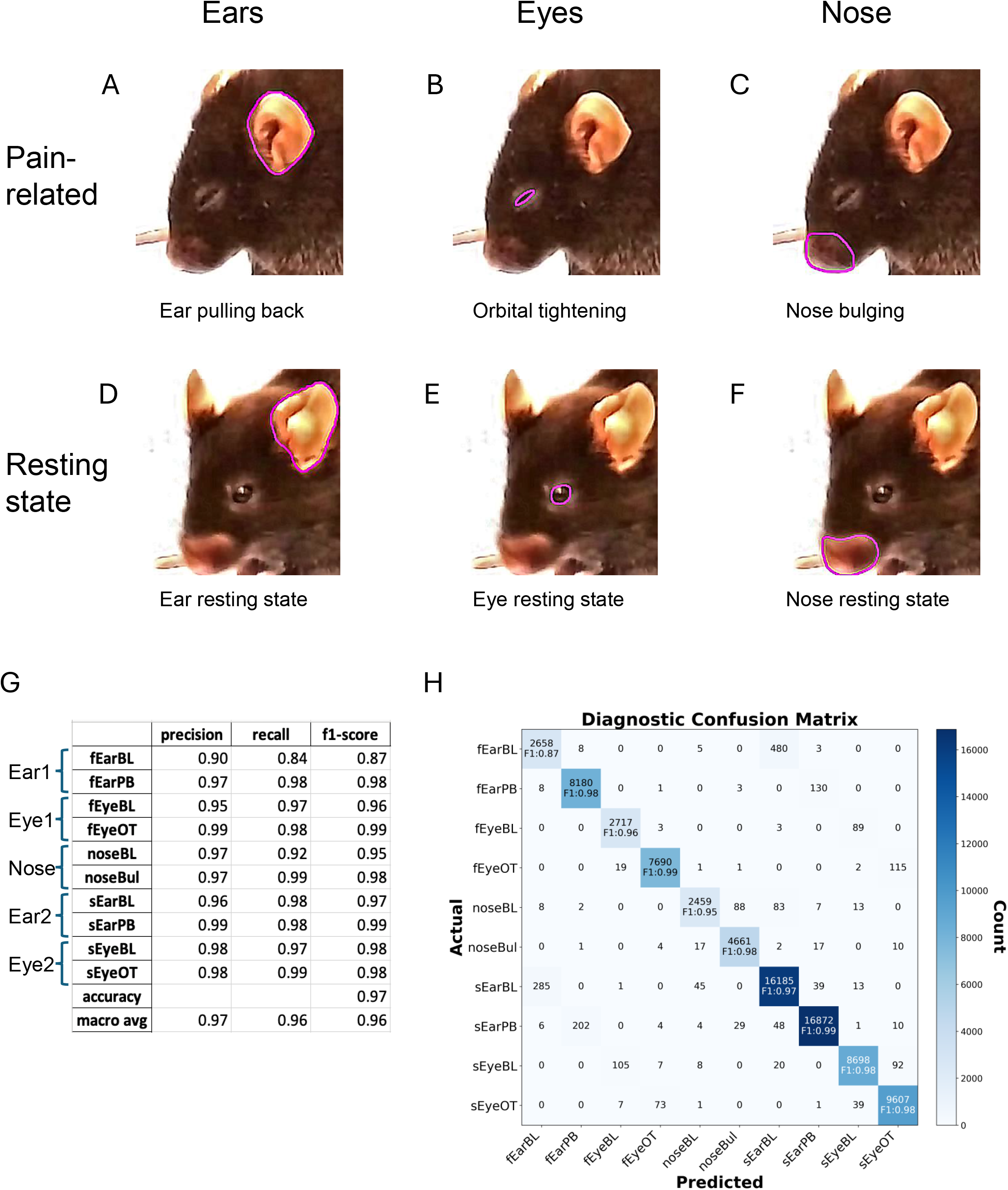
LabGym accurately analyzes pain-related facial action units in black-coated mice. Pink outlines indicate the facial regions associated with each facial feature state. **(A)** Ear position in the pain-related state. The ears are pulled apart and backward from their baseline position, often accompanied by vertical ridges formed as the ear tips retract. **(B)** Orbital tightening in the pain-related state. The orbital area is narrowed, characterized by a tightly closed eyelid or eye squeezing, denoted by wrinkles around the eye. **(C)** Nose bulge in the pain-related state. A rounded extension of skin is visible along the bridge of the nose. **(D)** Ear position in the resting state. The ears remain in their natural baseline position without retraction or ridge formation. **(E)** Eye position in the resting state. The orbital area shows no tightening, eyelid closure, or wrinkling. **(F)** Nose shape in the resting state. No rounded extension of skin is visible on the nasal bridge. **(G)** Per-class precision and overall classification performance of the trained facial-feature classifier across ten categories. The classifier achieved an overall accuracy (f1-score) of 0.97 and macro-averaged precision of 0.96 in f1-score. **(H)** Confusion matrix depicting classification performance across ten categories, including frontal (f) and side (s) views of ear baseline (EarBL), ear pulling back (EarPB), eye baseline (EyeBL), orbital tightening (EyeOT), nose baseline (noseBL), and nose bulge (noseBul). Diagonal entries show correctly classified samples and the F1 score for each category, whereas off-diagonal entries represent misclassifications. Color intensity reflects the proportion of samples in each predicted category. For downstream pain scoring, frontal-and side-view predictions for each facial region (ears, eyes, and nose) are combined to generate a single pain-related behavior score for that region.

To further evaluate the Categorizer’s performance, we manually annotated 500-second videos of mice with or without intraperitoneal administration of MgSO₄, a model of peritoneal irritation (Mogil et al. 1999) that has been shown to induce grimace in mice (Langford et al. 2010). These videos were used to generate an independent testing dataset that was not used in training the Categorizer. We then compared these annotations with the classifications produced by LabGym (Supplementary Figure 2). On this independent testing dataset, the Categorizer maintained a high overall weighted-average F1 score of 0.86. Performance varied across classes, with per-class scores ranging from 0.26 to 1.00. The lowest scores were observed for near-threshold or visually ambiguous classes (fEarPB, 0.26, as ear pulling back is barely visible frontally; noseBL, 0.41, as the resting nose resembles mild bulging), whereas the highest scores were observed in visually distinct classes (fEyeOT, 1.00; fEyeBL, 0.98; sEyeOT, 0.98). These results suggest that the remaining classification errors were primarily driven by ambiguity in borderline facial action units rather than by limitations of the model itself.

### A subset of kinematic descriptors correlate with pain levels across all three facial regions

A graded pain metric is advantageous over a binary pain metric because it can provide information on the relative level of pain an animal might be experiencing. To create a graded pain metric, we first generated a dataset encompassing multiple pain levels in mice through chemogenetic activation of *Scn10a*+ neurons, which comprise a diverse group of nociceptors (Mouchbahani-Constance et al. 2023). We reasoned that we could use a chemogenetic approach to control the relative level of nociceptive input by injecting Scn10a-Cre CAG-LSL-Gq-DREADD mice with different doses of the synthetic ligand, clozapine-N oxide (CNO) (Ronan et al. 2025). In this experiment, mice were recorded under six different conditions: an unperturbed true baseline group that received no injection, a baseline group that received a vehicle (phosphate buffered saline, PBS) injection, and groups that received one of the four CNO doses (0.01 mg/kg, 0.1 mg/kg, 0.5 mg/kg, and 1.0 mg/kg). Each condition used an experimentally naïve mouse, except for the vehicle and 0.5 mg/kg conditions where mice underwent a vehicle trial days before the 0.5 mg/kg CNO trial.

While the current version of LabGym outputs 13 quantitative measurements, only 9 are kinematic measurements that reflect the temporal changes of animals’ facial changes. Among the nine measurements contributed by each facial action unit, many would be expected to covary. Retaining all 27 features (9 features/region x 3 regions) therefore would add redundancy without improving prediction. To reduce this redundancy, we examined facial-region-specific pairwise Pearson correlation matrices across the CNO concentrations together with PBS and true baseline recordings (Figure 3A, C, E). When two measurements were correlated at r > 0.7, a threshold above which collinearity substantially distorts model estimation (Dormann et al. 2013), we retained the measurement with the larger across-condition variance (Figure 3B, D, F). Measurements with low variance across CNO conditions were subsequently excluded, as they demonstrated limited responsiveness to nociceptive stimulation.

**Fig. 3.**
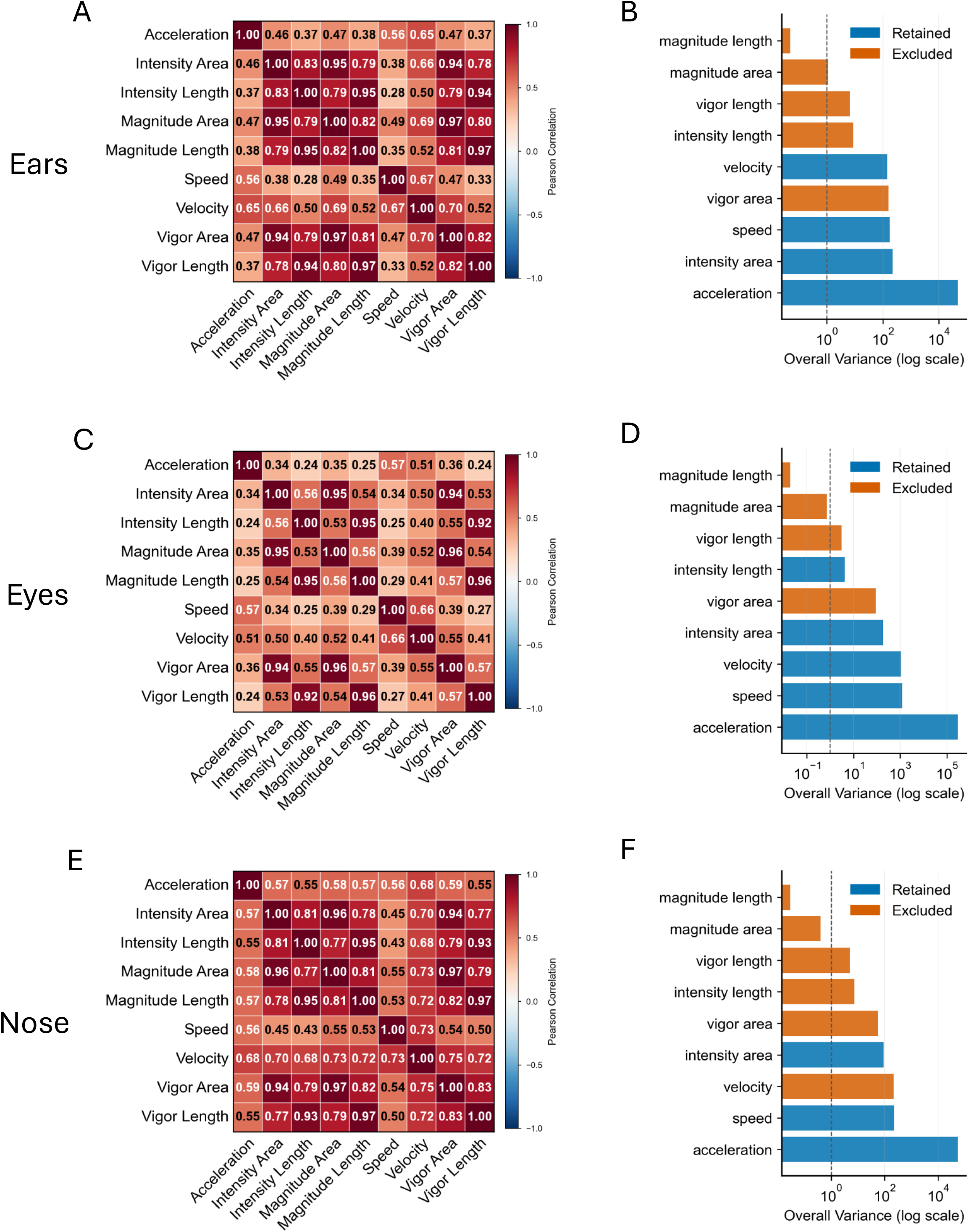
Kinematic descriptors across ear, eye, and nose facial action units correlate with pain levels. **(A, B)** Pearson correlation matrix **(A)** and feature-variance plot **(B)** for ear-position measurements. Pairwise features with a correlation coefficient r > 0.7 were considered redundant. For each correlated pair, the feature with the higher variance feature was retained (blue), whereas the lower-variance features was excluded (orange). Four features were retained. **(C, D)** Pearson correlation matrix **(C)** and feature-variance plot **(D)** for eye orbital-tightening measurements. The same selection criteria were applied, resulting in the retention of five features. **(E, F)** Pearson correlation matrix **(E)** and feature-variance plot (**F**) for nose-bulge measurements. The same selection criteria were applied, resulting in the retention of three features. The dashed vertical lines in pane **B-F** indicate the minimum variance threshold (1.0).

From the selected measurements, kinematic descriptors that were common among the three facial action units were retained. Acceleration, intensity area, intensity length, speed, and velocity were retained for both eye orbital tightening and nose bulging (Figure 3B, D, F). For ear position, intensity length was excluded during the variance-selection step.

Among the retained measurements, *intensity area* showed the strongest monotonic relationship with CNO concentration (ear resting state: r =-0.993; eye orbital tightening: r =-0.987; and nose bulging: r =-0.932) (Figure 4A) and exhibited the clearest dose-dependent trend when CNO concentration was plotted on a logarithmic scale (Figure 4B, C, D). Intensity area represents cumulative proportional change in a facial region across consecutive frames (n) within a time window, thereby capturing the overall area (a) change of the facial region over time (t) (Hu et al. 2023). Here, a diminished intensity area reflects sustained tonic muscle contractions (Kunz et al. 2019) and an extended expression of a given facial shape.

**Fig. 4.**
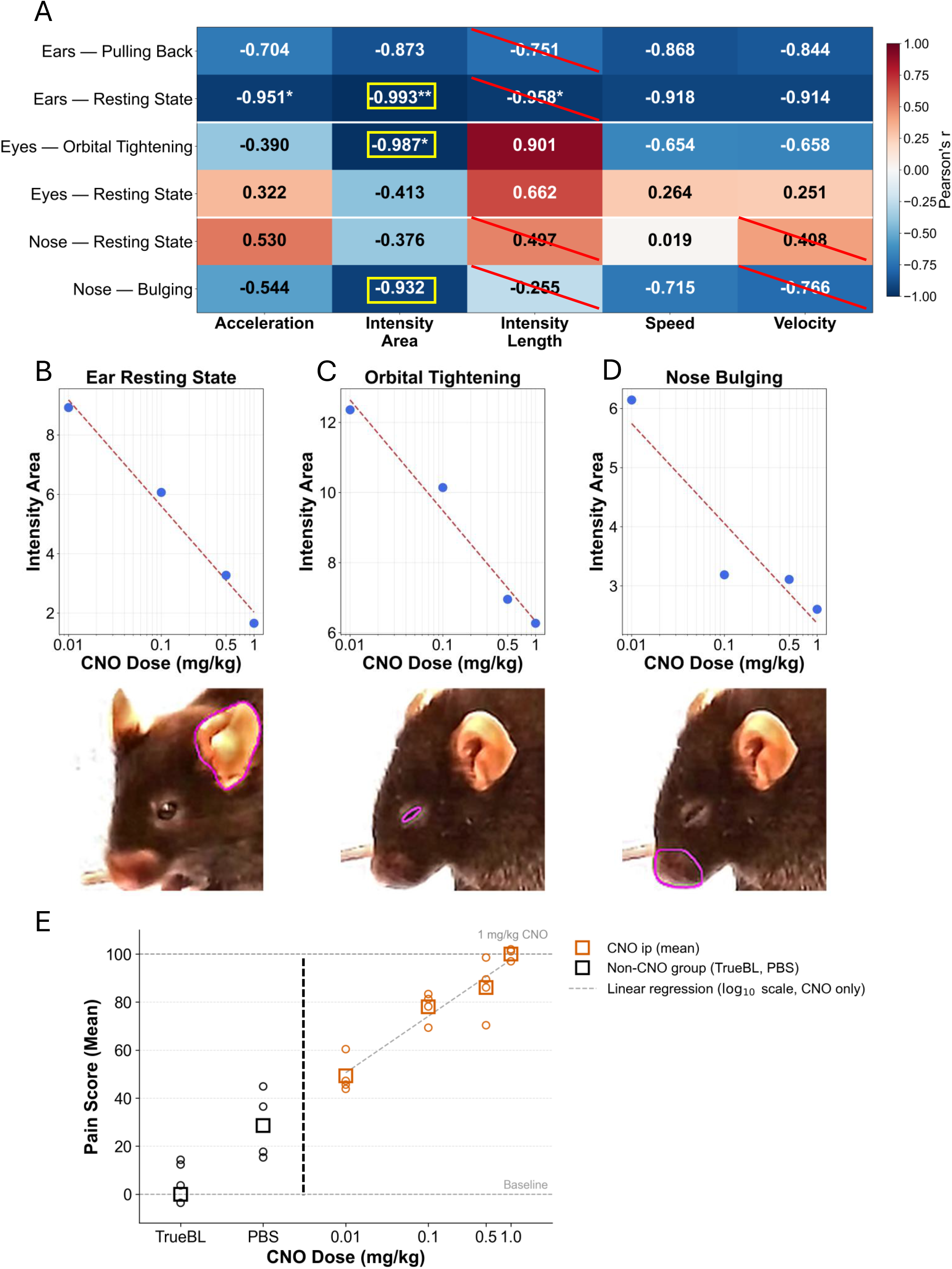
Quantification of kinematic descriptor correlations with pain levels for pain scoring. (A) Pearson correlation heatmap of facial features and log(CNO dose) for ears, eyes, nose regions. Highlighted cells show features with the strongest correlation to CNO dose. Features used for composite pain score are ears resting state, eye orbital tightening, and nose bulging (intensity area). **(B–D)** Linear regression of intensity area and log(CNO dose) for each selected facial regions. The slope of each regression is used as feature weight in composite pain score. Mouse images are shown below **(B)** ears resting state, **(C)** eye orbital tightening and **(D)** nose bulging. **(E)** Composite pain score as a function of CNO dose (mean SEM; n = 24, log scale). Pain score is calibrated to 0 = true baseline and 100 = 1.0 mg/kg CNO.

### LabGrymace produces a graded pain score from weighted facial action units

The dose-dependent relationships between CNO concentration and the kinematic measurement *intensity_area* also enabled us to determine how strongly each facial action unit contributed to the pain-related facial expression. Facial action units with steeper dose-response relationships to nociceptor activation received greater weight in the composite score. Accordingly, the slopes of corresponding dose-response curves were used to derive the weight of each facial action unit: W_EAR = 3.574, W_EYE = 3.158, and W_NOSE = 1.689 (Figure 3B-D).

Conceptually, the pain score was calculated in three steps. First, the measurement from each visible facial region was normalized relative to its CNO reference distribution so that signals from different regions could be compared on a common scale. Second, the normalized values were combined using the region-specific weights, with more pain-responsive facial action units contributing more strongly to the composite score. Third, the composite value was linearly calibrated such that the uninjected baseline corresponded to 0 and the 1.0 mg/kg CNO condition corresponded to 100. Thus, the final score provides a continuous measure of the strength of the pain-related facial response relative to these biological reference points.

The corresponding calculations were:

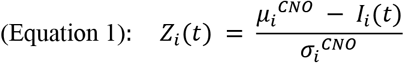

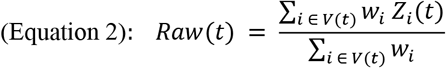

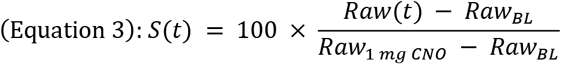

In Equations 1-3, *I_i_*(*t*) denotes the trimmed rolling-window intensity-area signal for the facial region *i* at time *t*; *μ_i_^CNO^* and *μ_i_^CNO^* denote the corresponding CNO reference mean and standard deviation; *V*(*t*) denotes the set of visible facial regions at time *t*; *Raw*(*t*) denotes the weighted composite before rescaling; *Raw_BL_* and *Raw*_1*mgCNO*_ denote the baseline and 1.0 mg/kg CNO calibration anchors; and *S*(*t*) denotes the final pain score.

Under this calibration scheme, the composite pain score increased monotonically across the CNO series (Pearson r = 0.920, p < 0.001, n = 24; Figure 4E). When CNO dose was displayed on a logarithmic scale, the score increased linearly across the tested range. This dose-response relationship was consistent with the Weber-Fechner law, which posits that perceived intensity scales with the logarithm of stimulus intensity (Boring 1961, Fechner et al. 1966). In this setting, CNO dose serves as the ordered stimulus input, whereas the LabGrymace composite serves as a behavioral proxy for internal nociceptive intensity. The score was calibrated across graded nociceptor activation and remained consistent with established stimulus-response principles. This represents a key distinction from static, appearance-based frameworks such as MGS and PainFace (Langford et al. 2010, McCoy et al. 2024), which are designed to reproduce human-assigned grimace scores.

### LabGrymace and PainFace employ distinct temporal strategies for facial action unit analysis

As a benchmark, we implemented and compared two distinct behavioral analysis pipelines to assess pain-related facial expression in mice: LabGrymace and PainFace. LabGrymace employs a rolling window approach in which consecutive frames are grouped into overlapping windows with a stride of 1 frame, allowing the system to capture the temporal dynamics of facial movement frame-by-frame (Figure 5A). For each window, the LabGrymace detector extracts three facial regions (i.e., ears, eyes, and nose) and generates a corresponding pattern image that encodes both the spatial trajectory and morphological changes of each facial region over the duration of the window (Figure 5B). The kinematic information embedded in these pattern images, including shape deformation and displacement, forms the basis for downstream behavioral classification.

**Fig. 5.**
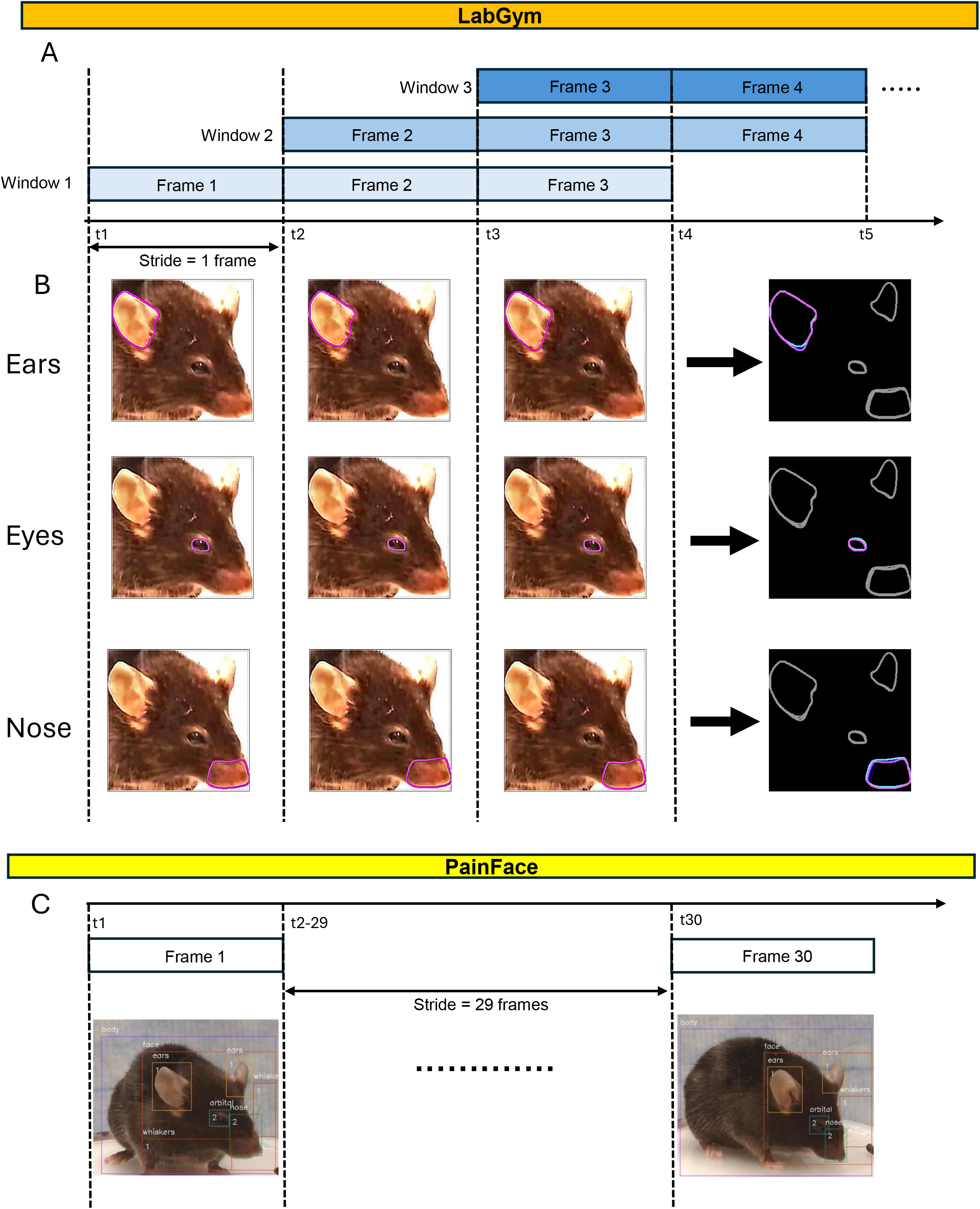
Comparison of behavioral analysis pipelines between LabGym and PainFace. **(A)** Schematic of the rolling window approach used in LabGym. Consecutive frames are grouped into windows of fixed size with a stride of 1, such that each window overlaps with the previous by all but one frame. **(B)** Ear region extracted by the LabGym detector. Left panels show the cropped animation frames at successive time points (t1-t3), with the ear region highlighted with pink outlines. Right panel shows the corresponding pattern image generated from the rolling window, in which the spatial trajectory and morphological changes of the focal facial regions across the window are captured; the blue and magenta contours represent the focal facial regions at the first and last frames of the window, respectively, illustrating its kinematics over time, while gray contours represent other non-focal facial regions detected within the same frame. **(C)** Eye regions extracted by the LabGym detector, displayed in the same format as (B). **(D)** Nose region extracted by the LabGym detector, displayed in the same format as (B). **(E)** Representative output of the PainFace pipeline. Input video is sampled at 1 frame per second, and each sampled frame is independently evaluated for four facial action units. Only frames in which all four facial action units are successfully detected are retained. Shown are the first (Frame 1) and last (Frame 30) retained frames with detected facial action units annotations overlaid.

PainFace adopts a fundamentally different approach by treating each frame as an independent unit of analysis (McCoy et al. 2024). Input video is sampled at 1 frame per second, and each sampled frame is evaluated for four facial action units individually (Figure 5C). Only frames in which all four facial action units are successfully detected are retained for scoring, after which pain scores are averaged across valid frames within a defined time window. Unlike LabGym, PainFace does not model inter-frame dynamics or generate pattern images; instead, it relies on single-frame detection to quantify facial action unit expression. These two pipelines thus differ fundamentally in their temporal resolution, feature representation, and sensitivity to moment-to-moment changes in facial expression.

### LabGrymace quantifies the intensity and temporal dynamics of visceral pain

We next asked whether LabGrymace could reliably quantify pain in an established model for pain. Intraperitoneal administration of MgSO₄ is a standard model of peritoneal irritation (Mogil et al. 1999) that has previously been shown to evoke grimace in mice (Langford et al. 2010). We therefore used this visceral pain model to determine whether LabGrymace could detect pain-related facial changes already recognized within the MGS framework, while also testing whether the calibrated pain score generalized across pain modality and stimulus intensities.

To this end, we tested a range of MgSO₄ doses,including 62.5, 125, and 250 mg/kg, encompassing the dose known to reliably elicit grimace (Langford et al. 2010). As expected, 125 and 250 mg/kg resulted in robust pain scores throughout the 500-s evaluation (Figure 6A, B). However, LabGrymace-derived pain scores revealed additional unexpected insights into dose-dependent pain dynamics across both dose conditions and individual 100-s time windows.

**Fig. 6.**
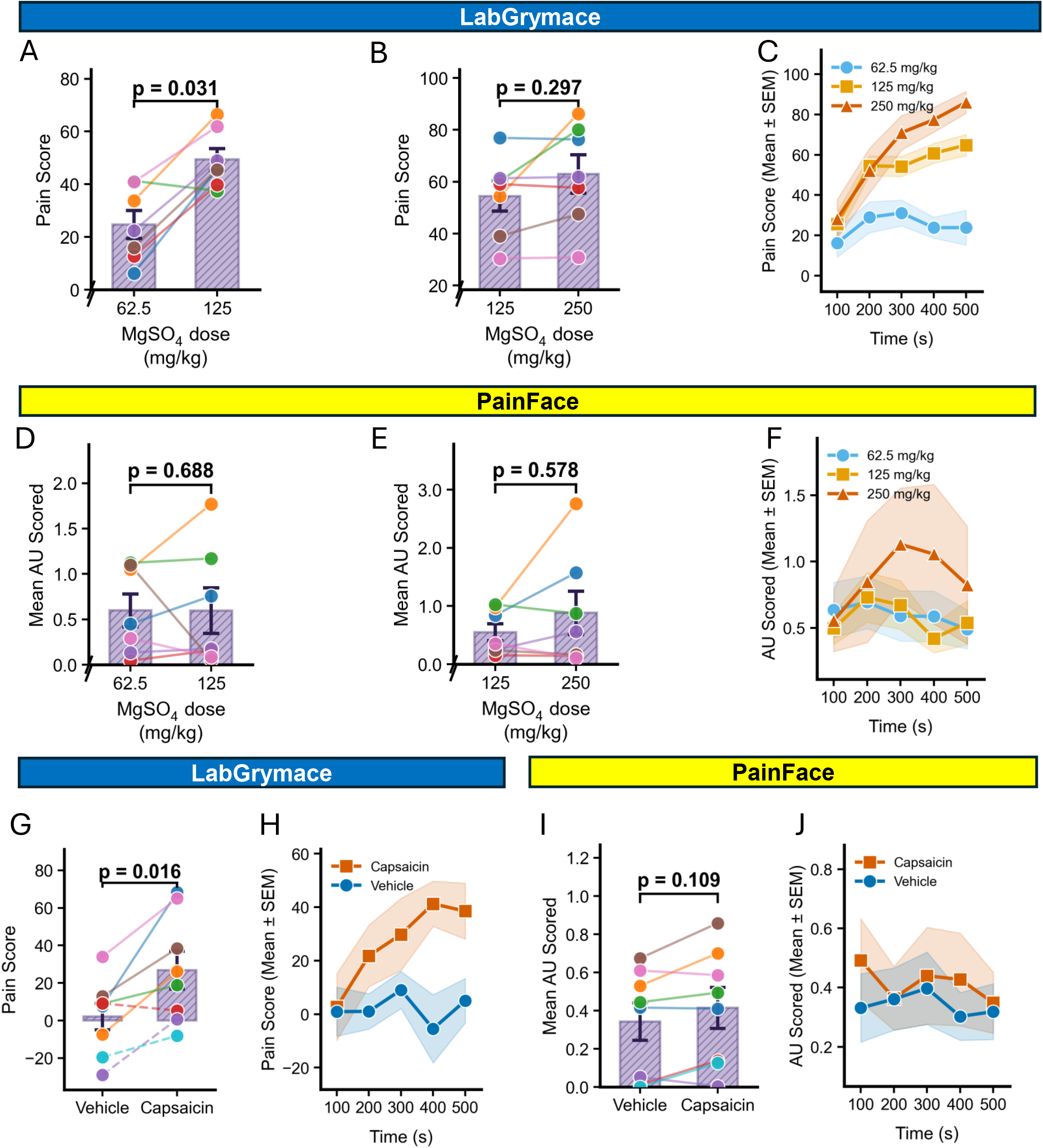
LabGrymace quantifies the intensity and temporal dynamics of pain in visceral (MgSO_4_) and somatic (capsaicin) pain models. Each time window represents 100 seconds (3000 frames acquired at 30 frames/second). **(A–C)** LabGrymace pain scores in mice treated with MgSO_4_. (**A**) 62.5 and 125 mg/kg MgSO_4_. (B) 125 and 250 mg/kg MgSO_4_. (**C**) Time course of pain scores for all three doses (62.5, 125, and 250 mg/kg; mean ± SEM). Mean LabGrymace pain scores increased with MgSO_4_ dose. N = 7 in all groups. **(D–F)** PainFace mean facial action unit (AU) scores in MgSO_4_-treated mice. (**D**) Mean AU scores at 62.5 and 125 mg/kg MgSO_4_. (**E**) Mean AU scores at 125 and 250 mg/kg MgSO_4_. (**F**) Time course of mean AU scores for all three doses (mean ± SEM). **(G–H)** LabGrymace pain scores for capsaicin-treated mice. (**G**) vehicle and capsaicin (125 µg/mL). (**H**) Time course of pain scores for vehicle and capsaicin groups (mean ± SEM). Capsaicin treatment produced significantly higher pain scores than vehicle controls. N = 8. **(I–J)** PainFace mean AU scores for capsaicin-treated mice. (**I**) Mean AU scores for vehicle and capsaicin (125 µg/mL) groups. (**J**) Time course of mean AU scores for vehicle and capsaicin groups (mean ± SEM). All paired comparisons were assessed by Wilcoxon signed-rank tests.

Importantly, LabGrymace generated distinct pain score dynamics for each of the three MgSO₄ doses tested. Evaluation of consecutive 100-s windows revealed that all doses elicited increasing pain scores during the initial two 100-s windows (Figure 6C). However, only the higher doses (125 and 250 mg/kg) produced increasing pain scores across all time windows. The step from 62.5 to 125mg/kg produced a significant increase in pain score (p = 0.031, n = 7, Figure 6A), whereas the step from 125 to 250 mg/kg raised the mean score but did not reach significance at the current sample size (p = 0.297, n = 7, Figure 6B). Notably, pain scores for the two highest doses diverged during the later 100-s windows (300-500 s). Although the 250 mg/kg group trended higher than 125 mg/kg at the later windows, the two doses were not statistically separable over the 500-s recording. This may indicate either that their trajectories diverge only beyond the analyzed 500-s window, or that the response at 125 mg/kg already approaches a plateau, limiting further dose-dependent separation at these higher doses.

Across all 28 recordings, pain scores showed a positive overall association with MgSO4 dose (Pearson r = 0.608, p < 0.001), indicating that the CNO-calibrated scale remained ordered when applied to a visceral pain model.

To compare performance with an existing automated approach, we applied PainFace (McCoy et al. 2024) to the same video recordings. Although PainFace identified a trend toward increased pain with increasing MgSO₄ dose, dose-dependent differences were not appreciable at our current sample sizes. Moreover, action-unit scores generated by PainFace were highly variable across both time and dose conditions (Figure 6D-F), obscuring differences in pain-response kinetics.

Taken together, these findings demonstrate that LabGrymace and its calibrated pain scale can detect pain-related facial changes in an established model of visceral pain, while revealing new insights into the intensity and temporal dynamics of pain induced by varying doses of an irritant.

### LabGrymace quantifies the distinct temporal dynamics of capsaicin-induced somatic pain

Given the sensitivity of LabGrymace in detecting pain at the lowest dose of MgSO_4_, we next asked whether it could identify pain-associated facial features in an empirically variable model of induced pain. Capsaicin is one of the earliest and most widely used chemical stimuli for activating nociceptors via agonism of TRPV1 (Caterina et al. 1997, Mogil et al. 2006). However, capsaicin injection initially failed to evoke reproducible facial grimace in mice (Langford et al. 2010). We reasoned that the continuous sampling and quantitative pain scoring by LabGrymace might provide improved resolution for this model of acute chemically induced pain.

To test this, we administered a single dose of capsaicin (125 μg/ml) via intraplantar injection to replicate prior studies (Langford et al. 2010). In line with the sensitivity of LabGrymace, we found elevated pain scores following capsaicin injection compared to vehicle-treated group (Figure 6G). As would be expected with prior variability in detecting capsaicin-induced facial grimace, the pain scores were at the lower end of our scale and were comparable to those elicited by the lowest dose of MgSO_4_.

Analysis of consecutive 100-s time windows further revealed a temporal profile of capsaicin-induced pain that was distinct from that observed flowing MgSO_4_ administration. Specifically, pain scores began to increase during the second 100-s window after capsaicin injection, and continued to increase and then plateaued during subsequent intervals.

As with MgSO_4_ injections, PainFace identified a trend of action-unit score increase following capsaicin injection; however, these changes were not significantly different from vehicle controls at the sample sizes evaluated. In addition, PainFace action-unit scores exhibited marked variability across time within each condition, yielding temporal profiles that were incongruous with both LabGrymace-derived pain scores and previously reported pain kinetics following capsaicin injection measured using alternative behavioral assays (Frias et al. 2016).

Taken together, these finding demonstrate the sensitivity and versatility of LabGrymace for identifying and quantifying facial grimace associated with acute somatic pain while providing enhanced temporal resolution of pain dynamics.

## DISCUSSION

Facial expressions are widely used by mammals to facilitate non-verbal communication as a possible indication of distress (Mota-Rojas et al. 2025). Grimace-based scoring systems have demonstrated that facial features can be leveraged to quantify pain-related behavior in laboratory animals (Langford et al. 2010, Sotocinal et al. 2011, Matsumiya et al. 2012, Hager et al. 2017). In this study, we establish LabGrymace, a customizable AI-based framework that detects discrete facial action units and generates quantitative pain scores in sub-second temporal resolution. This platform (1) builds on LabGym-based segmentation, feature detection, and kinematic quantification to track and measure facial function units in freely moving mice; (2) provides an open, transparent, and customizable workflow that can be readily adapted by other users; (3) quantifies pain-induced facial dynamics on a frame-by-frame basis and thus captures grimace dynamics with high temporal resolution; (4) assigns differential weights to facial action units based on their relationships with graded nociceptor activation; (5) integrates the weighted features into a composite pain score capable of detecting subtle pain-related facial changes; and (6) establishes a generalizable strategy that can be extended to identify facial or bodily features associated with other internal states.

While the trained Detector, Categorizer, and pain-score calculation parameters presented here are ready to use with acquisition parameters similar to those used in this study, they can be readily retrained and adapted to alternative imaging setups through the user-friendly LabGym and LabGrymace GUIs. Thus, our work provides a general framework for developing weighted, feature-based classifiers that can be extended to additional pain models, different animal species, and other internal or affective states reflected in facial movements.

The pioneering work defining the MGS demonstrated the utility of facial features in evaluating nociception and pain but also reported variable performance in identifying some forms of acute pain (Langford et al. 2010). Our findings indicate that pain-related facial changes may simply fail to be captured by conventional scoring approaches. The ability of LabGrymace to detect chemical nociception and the corresponding subtle and transient facial changes is a critical advance. By measuring multiple facial features and weighing them according to their relationship with nociceptive input, LabGrymace was able to identify low MgSO_4_-dose and capsaicin-related pain responses that cannot be detected using coarse manual scoring and a consolidated observation period (Langford et al. 2010). Furthermore, LabGrymace provided temporal dynamics of two models of acute pain, revealing critical differences in pain onset, apparent intensity, and duration. These data provide a more nuanced evaluation of pain and the basis for evaluating the efficacy and/or effects of analgesics. Our work simultaneously refocuses attention on facial behavior as a useful proxy of pain-related state and provides a basis for quantitative analysis of models of pain and its relief.

A key distinction between LabGrymace and prior facial pain-analysis approaches is its flexibility and sensitivity. PainFace demonstrated the value of automated facial analysis, but its implementation depends on uniform, prescribed acquisition conditions and a fixed analytical pipeline (McCoy et al. 2024). In contrast, LabGrymace is a customizable tool that allows users to define the relevant facial features, determine the extent to which those features covary with an experimental condition, and modify the analysis to their own imaging conditions or research questions. This flexibility is important because laboratory environments and animal appearance vary substantially across studies (Sadler et al. 2022). Moreover, PainFace analyzes only frames in which all facial action units are simultaneously visible, rendering frames with obscured features unsuitable for analysis. When sampled frames capture the mouse turned away from the camera, pain state cannot be inferred reliably from those images. In contrast, LabGrymace analyzes facial dynamics within short time-windows, evaluating kinematic changes on a frame-by-frame basis. This approach enables continuous inference of pain state from the visible facial regions even when some facial regions are occluded, resulting in more stable pain scores in drug-testing experiments in which changes in head orientation are unavoidable. These methodological differences likely contributed to the greater sensitivity of LabGrymace and its enhanced ability to detect differences in both pain intensity and kinetics at the sample sizes examined in this study.

A further advantage of this framework is that its quantitative output is grounded in an objective biological reference. Like all supervised grimace classifiers, the LabGym Categorizer is trained using manually annotated facial states, and such annotation inevitably contain a degree of subjectivity. Consistent with this, evaluation of the trained Categorizer on an independently annotated testing dataset yielded high overall accuracy but more variable performance in individual classes, particularly for the subtle and visually ambiguous action units (Supplementary Figure 2). This variability mirrors the inter-observer disagreement that has long constrained manual MGS scoring (Langford et al. 2010). Critically, however, LabGrymace is designed so that uncertainty at the labeling stage does not directly determine the final pain score. The Categorizer determines only which facial state is present, whereas the magnitude of the pain score is determined by action-unit-specific weights and calibration parameters derived from an objective, experimenter-controlled relationship—the CNO chemogenetic dose–response curve— rather than from human-assigned labels. Consequently, the resulting pain score depends on the underlying stimulus-response relationship captured by the model and is therefore less dependent on the classification accuracy of the LabGym Categorizer.

Anchoring the score to a graded, controlled stimulus further allows the framework to be validated against an external, theory-based prediction rather than against human ratings. Across the CNO dose series, the composite pain score increased monotonically and was approximately linear on a logarithmic dose scale (Pearson r = 0.920; Figure 4E), consistent with the Weber-Fechner law, which states that perceived intensity scales with the logarithm of stimulus intensity (Boring 1961, Fechner et al. 1966). Recovering this relationship from facial dynamics provides independent evidence that LabGrymace captures a genuine stimulus-response axis rather than merely reproducing subjective annotations. Thus, LabGrymace bridges a gap in automated grimace analysis: it converts facial movements into a quantitative pain readout whose validity rests on a well-established perceptual law, rather than on the consistency of manual scoring.

Our approach also provides insight into the generalizability of facial features as an indicator of pain-related behaviors. Nociceptive input did not appear to alter all facial features equally. Instead, a subset of features was more strongly associated with the intensity of pain, whereas other features contributed less or provided redundant information. This observation underscores the importance of weighting features rather than treating all features as equivalent indicators of pain. Thus, an open question is whether different pain modalities engage overlapping but non-identical facial features or identical facial features. Future studies using LabGrymace to compare pain models will be necessary to determine whether a common facial pain ‘axis’ exists or whether unique feature weighting is necessary for each pain modality.

We acknowledge inherent limitations of our study. Our work does not establish whether LabGrymace can detect spontaneous pain in preclinical models of chronic pain. The present experiments use graded nociceptive input based on chemogenetic activation of peripheral nociceptors as a standard. This design provides a clear basis for feature selection and score generation, but it does not directly address pain states in which animals are not externally stimulated closely preceding the time of measurement. Spontaneous pain remains challenging to define and measure in rodents, in part because animals may suppress overt signs of vulnerability (Tappe-Theodor et al. 2019). Future work will be needed to determine whether LabGrymace can identify ongoing pain-related facial states in models of inflammatory or chronic pain.

Moreover, our study has measured some, but not all possible facial features that could be associated with pain. The facial features identified here were selected based on their relationship to the nociceptive conditions examined in this study. It will also be important to determine which, if any, acquisition parameters might influence feature detection and pain score. These considerations are particularly important for the implementation of LabGrymace as a standardized tool across laboratories.

Despite these limitations, LabGrymace establishes a general framework for transforming dynamic facial behavior into quantitative measures of internal state. Its open, customizable design enables researchers to retrain the underlying models, define new behavioral features, and calibrate weighted scores for diverse experimental conditions, animal species, and behavioral paradigms. As AI-based behavioral analysis continues to evolve, frameworks that combine interpretable feature extraction with biologically grounded calibration may provide a general strategy for quantifying not only pain, but also a broad range of internal and affective states that are reflected in facial and body movements.

## MATERIALS AND METHODS

### Mouse strains and husbandry

All animal experiments followed NIH guidelines and were performed in accordance with protocols approved by the Institutional Animal Care and Use Committee at the University of Michigan. Mice were group-housed at room temperature and animal rooms were maintained on a 12-hour light/12-hour dark cycle. Mice had *ad libitum* access to standard lab mouse pellet food and water. Scn10a-Cre, CAG-LSL-Gq-DREADD (hM3Dq), and C57BL/6J mice were purchased from Jackson Laboratory. Female and male mice were used for all experiments.

### Behavioral apparatus and acclimation

Mice were contained in a plexiglass acrylic tube (3.78” inner diameter x 6” height) and were allowed to freely move within the tube. The tube was placed within a sound attenuating cubicle (interior: 25.5” width x 20.75” height x 21.5” depth; Med Associates, Inc.), and a webcam (Logitech C920S Pro 1080p) was placed approximately 8.5 inches away from the tube on a platform that was 3 inches tall. A rechargeable LED light source was placed above the tube. Blue absorbent lab soaker sheets were placed on the walls of the chamber to reduce glare and ensure a consistent background between videos. Before undergoing experimental treatments, mice underwent three days of acclimation. During acclimation, mice were placed in the behavioral chamber for 20 minutes. All acclimation and behavioral procedures occurred in the morning during the early hours of the light cycle in the animal housing rooms.

Although the Logitech webcam was configured to record at 30 fps, its effective frame rate fluctuated between 29 and 30 fps. Consequently, analyses assuming a nominal frame rate of 30 fps introduce a maximum timing error of approximately 3% over the 500-s integrated analyses shown in Figure 6. The resulting error is proportionally smaller for the 100-s temporal analyses in the same figure and does not affect the relative comparisons of pain scores across experimental conditions.

### Chemogenetics

The *Scn10a*-Cre driver was crossed into CAG-LSL-Gq-DREADD to enable selective activation of *Scn10a*+ sensory neurons. These neurons represent a large, diverse class of nociceptors and were activated using the chemogenetic drug clozapine-N-oxide (CNO). CNO was diluted in phosphate buffered saline (PBS) and administered at 0.01 mg/kg, 0.1 mg/kg, 0.5 mg/kg, and 1 mg/kg doses via intraperitoneal injection. A PBS-only vehicle control was included to account for pain-related behavioral changes associated with the injection procedure in the absence of chemogenetic activation of *Scn10a*+ neurons. An uninjected baseline group was also recorded to capture normal behavior in the absence of injection-induced pain. The PBS vehicle and 0.5 mg/kg treatments were performed on the same four mice, whereas the remaining conditions were tested in independent cohorts of mice (baseline, n = 5; 1.0 mg/kg CNO, n = 3; all other treatment groups, n = 4). Intraperitoneal injections were administered in awake, scruffed animals, which were subsequently placed in their home cage to rest for 20 minutes to allow sufficient time for CNO-mediated activation of *Scn10a*+ neurons. Following this 20-minute period, animals were placed in the behavioral chamber and 20-minute videos were immediately captured.

### Magnesium Sulfate & Capsaicin treatments

Mice with a C57BL/6J background were used for MgSO_4_ and capsaicin experiments. MgSO_4_ was diluted in saline and administered intraperitoneally at 62.5, 125, and 250 mg/kg doses. Two cohorts of mice received the MgSO_4_ treatment. One group received the 62.5 mg/kg dose and then received 125 mg/kg a few days later, while a second group received the 125 mg/kg dose and then received 250 mg/kg. Capsaicin was dissolved in a 10% ethanol, 10% Tween 80, and 80% saline solution. Mice received a 20 µL intraplanar injection of a 125 µg/mL capsaicin solution or a vehicle-only solution. For both MgSO_4_ and capsaicin experiments, mice were placed in the behavioral chamber, and 20-minute videos were captured immediately following injection.

### Computational hardware

Computations were performed on the spatially programmed GPU (SPGPU) partition of the University of Michigan Great Lakes cluster. The SPGPU nodes are equipped with two 2.9 GHz Intel Xeon Gold 6226R processors (32 cores per node), 376 GB RAM, and eight NVIDIA A40 GPUs with 48 GB memory each. Detector and Categorizer training, as well as subsequent computational analyses, were performed using this hardware configuration.

### Behavioral video processing

Behavioral videos were recorded at 30 frames/s for 20 min per session. For LabGrymace analysis, only the first 500s of each recording were used because this interval captured the most pronounced post-treatment responses while reducing unnecessary computational burden. Videos were cropped to 1080 x 1080 pixels so that the mouse remained centered in the field of view, and contrast was uniformly adjusted with a factor of 2 to preserve visibility of the eyes. Occasional reflections from the chamber wall or transient occlusion by fur could reduce eye visibility, although such cases were infrequent. These preprocessed videos were then used for downstream Detector, Categorizer, and measurement-based analyses in LabGym and LabGrymace.

### The architecture of neural network

#### Detector

The Detector in LabGym was based on Mask R-CNN with a ResNet-50 backbone and was used to detect and segment the facial regions of interest before downstream classification and measurement extraction.

#### Animation Analyzer

The Animation Analyzer extracts temporal information from Animation examples using time-distributed convolutional layers followed by long short-term memory (LSTM) layers. LabGym provides seven complexity levels spanning VGG-like and ResNet-like architectures. In the present study, the Animation Analyzer was run at complexity level 3 (LV3).

#### Pattern Recognizer

The Pattern Recognizer extracts spatial information from the Pattern Image using convolutional layers operating on RGB images that encode the temporal sequence of movements. As in the Animation Analyzer, LabGym provides seven VGG-like and ResNet-like complexity levels. Unlike the Animation Analyzer, the Pattern Recognizer processes static pattern images rather than time-distributed inputs. In the present study, the Pattern Recognizer was also run at LV3.

#### Decision Maker

The Decision Maker concatenates the outputs of the Animation Analyzer and Pattern Recognizer and passes the merged representation through fully connected dense layers to generate the final behavioral classification probabilities.

Together, these modules allow LabGym to combine temporal facial dynamics with static pattern-based features in a single categorization framework. Additional architectural details, including normalization and regularization layers, have been described previously (Hu et al. 2023).

### Detector training in *LabGym*

The LabGym Detector, *2026.4.6 detector Facial expression*, adapted from Detectron2 and described previously (Goss et al. 2024), was trained and used in this study to detect and segment five facial regions of interest in freely moving mice: the left ear, right ear, left eye, right eye, and nose.

Image examples were first generated from facial videos using the “Generate Image Examples” feature in the Train Detectors submodule of LabGym. A total of 496 images were manually annotated in Roboflow as five facial region classes corresponding to the left ear, right ear, left eye, right eye, and nose. The annotated images were then augmented in EZannot, which generated 134 additional manipulated images from each original annotated image, yielding a 135-fold image set for detector training in LabGym.

Detector training was performed with an inferencing frame size of ranging from 550-1400 px in width and 700-930 px in height, depending on the position of the mouse, for 10,000 training iterations. Frame size showed no correlation with pain score once batch was accounted for (within-batch Pearson r =-0.015, p = 0.963). Detector performance was evaluated using COCO-style metrics for bounding-box detection and instance segmentation. Overall, the Detector achieved a bounding-box mean average precision (mAP) of 89.4% and a segmentation mAP of 79.0%. For bounding-box detection, per-category AP was 94.7% for ears, 81.5% for eyes, and 92.1% for nose. For instance segmentation, per-category AP was 89.0% for ears, 63.5% for eyes, and 84.3% for nose. Performance was therefore lowest for the eye region, particularly in instance segmentation, consistent with the smaller size and more variable visual boundaries of the eyes in freely moving mice.

### Categorizer training in LabGym

The Categorizer, *LabGrymace_categorizer*, trained in LabGym was used to classify facial action units in LabGrymace. The Categorizer was trained using the Train Categorizers submodule embedded within the Training Module of LabGym. Training examples were augmented using the five default methods built into the LabGym, including random rotation, horizontal flipping, vertical flipping, random brightening, and random dimming. The Categorizer incorporated both the Animation Analyzer (complexity level 3) and Pattern Recognizer (complexity level 3) submodules and was trained on behavior examples generated in the Advanced-interactive mode. Behavior examples were generated using a 3-frame sliding window with a stride of 1 frame, such that each window yielded one Animation and its paired Pattern Image. Using the Analyze Behaviors module, predictions were updated frame by frame. The retained training settings were Animation Analyzer (LV3, 16 x 16 x 3) and Pattern Recognizer (LV3, 16 x 16 x 3). We manually sorted 8,619 training and 2,155 (80%: 20% split) testing behavior examples across 10 facial-state classes defined over five tracked facial regions: the left ear, right ear, left eye, right eye, and nose. These classes comprised frontal and side views of ear baseline (fEarBL, sEarBL), ear pulling back (fEarPB, sEarPB), eye baseline (fEyeBL, sEyeBL), orbital tightening (fEyeOT, sEyeOT), nose baseline (noseBL), and nose bulging (noseBul). Categorized examples were sampled from 5-min baseline and 0.1 mg/kg CNO videos, enriched for less frequent or harder-to-classify facial states, and split 80%/20% into training and validation sets. Both the training and validation sets were expanded 38-fold (see data-augmentation methods details below) by the default LabGym augmentation pipeline (327,522 and 81,890 examples). By contrast, the independent testing dataset, comprising 967 manually labeled examples from MgSO₄ videos, was not augmented.

The hyperparameters used for training the Categorizer have been described previously (Hu et al., 2023). LabGym uses an automatic training pipeline that determines appropriate training settings based on dataset size and training progress. Training was performed with stochastic gradient descent and cross-entropy loss, and the learning rate and model updates were adjusted according to validation performance as described previously. In the present study, Categorizer training in LabGym used its default batch size of 32.

The data-augmentation methods are as follows. Under the default LabGym augmentation scheme used here, each original behavior example was expanded into 38 aligned variants.

### Random rotation

Rotation augmentation included six preset angle conditions and one additional random-rotation condition. The six preset conditions corresponded to rotations sampled from 5°-45°, 45°-85°, and 95°-135° and 135°-175°, together with fixed 90° and 180° rotations.

### Horizontal flipping

Animations and their paired pattern images were flipped left-right. Horizontal flipping could also be combined with other default augmentation operations.

### Vertical flipping

Animations and their paired pattern images were flipped top-bottom. Vertical flipping could be applied alone or jointly with horizontal flipping and other default augmentation operations.

### Random brightening

Brightness was randomly increased within the default LabGym brightening range, corresponding to positive intensity offsets applied to the image.

### Random dimming

Brightness was randomly decreased within the default LabGym dimming range, corresponding to negative intensity offsets applied to the image.

Under the default LabGym augmentation scheme used here, each original behavior example was expanded 38-fold. This expansion arose from the original example, six preset rotation variants, and additional combinations of random rotation, horizontal flipping, vertical flipping, random brightening, and random dimming. Identical transformations were applied to both the animation and its paired pattern image to preserve alignment during training.

### Criteria for sorting the behavior examples

To train the Categorizer, we sorted behavior examples generated in the Advanced-interactive mode of LabGym into facial-action categories. Examples were drawn from 5-min baseline and 5-min 0.1 mg/kg CNO videos from two mice, with the two conditions contributing approximately equally to the categorized examples. Labels were assigned strictly according to MGS-defined facial morphology rather than treatment group. Thus, if an image or behavior example from the 0.1 mg/kg CNO condition still displayed baseline-like ear, eye, or nose morphology, it was labeled as the corresponding baseline class. This design ensured that the Categorizer learned objective MGS-based facial states rather than treatment labels, thereby informing downstream measurement selection and feature weighting without directly determining the final pain score. The per-class counts reported below correspond to the unaugmented dataset used for internal categorizer training (Figure 1D-E). One example pair consisted of one Animation and its corresponding Pattern Image.

### Ears

Frontal ear baseline (fEarBL; 416 examples): baseline-like ear position in which the ear remained in its resting configuration.

Frontal ear pulling back (fEarPB; 1094 examples): pain-related state in which the ear was retracted posteriorly in a manner consistent with the MGS.

Side-view ear baseline (sEarBL; 2182 examples): baseline-like ear position viewed laterally.

Side-view ear pulling back (sEarPB; 2258 examples): pain-related lateral-view state in which the ear was retracted posteriorly according to the same MGS-based criterion.

### Eyes

Frontal eye baseline (fEyeBL; 372 examples): baseline-like open eye without evidence of orbital tightening.

Frontal orbital tightening (fEyeOT; 1028 examples): narrowing of the orbital area with a tightly closed eyelid or eye squeeze, consistent with the MGS.

Side-view eye baseline (sEyeBL; 1175 examples): baseline-like open eye viewed laterally.

Side-view orbital tightening (sEyeOT; 1281 examples): lateral-view orbital narrowing defined using the same MGS-based criterion.

### Nose

Nose baseline (noseBL; 350 examples): baseline-like nose morphology without visible bulging.

Nose bulging (noseBul; 618 examples): rounded extension of skin visible on the bridge of the nose, defined according to the MGS.

Together, these criteria yielded 10 facial-state classes defined over five tracked facial regions: the left ear, right ear, left eye, right eye, and nose. Class labels were assigned according to visible facial morphology rather than treatment condition.

### Categorizer Testing

Categorizer performance was internally evaluated in LabGym using the Test Categorizers function with a validation dataset containing the same 10 facial-state categories used for training, but composed of behavior examples excluded from the training dataset. The validation data were augmented using the same five default augmentation methods used during training.

The augmented validation dataset used for internal testing contained the following category counts:

*frontal ear baseline* (fEarBL), 3,154 examples;

*frontal ear pulling back* (fEarPB), 8,322 examples;

*frontal eye baseline* (fEyeBL), 2,812 examples;

*frontal orbital tightening* (fEyeOT), 7,828 examples;

*nose baseline* (noseBL), 2,660 examples;

*nose bulging* (noseBul), 4,712 examples;

*side-view ear baseline* (sEarBL), 16,568 examples;

*side-view ear pulling back* (sEarPB), 17,176 examples;

*side-view eye baseline* (sEyeBL), 8,930 examples;

*side-view orbital tightening* (sEyeOT), 9,728 examples.

The following prediction equations were used to calculate performance metrics of Categorizers, including precision (Equation 1), recall (Equation 2), F1 score (Equation 3), and overall accuracy (Equation 4).

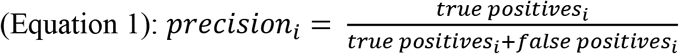

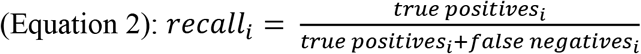

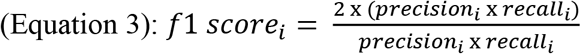

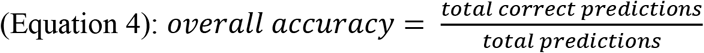

Confusion matrix (Figure 2H): The confusion matrix was generated using the scikit-learn library in Python 3.10 from the validation dataset used for internal categorizer testing. Predicted labels were generated by the LabGrymace categorizer, and true labels were derived from the corresponding ground-truth validation annotations for the same 10 facial-state categories.

### LabGrymace analysis

#### Summary file generation

LabGrymace analysis began with the processed output generated by the Detector and Categorizer modules of LabGym (Figure 1H). Measurement files corresponding to baseline and pain-related ear, eye, and nose states were consolidated into three region-specific summary files (ear_summary.xlsx, eye_summary.xlsx, and nose_summary.xlsx), each containing frame-level measurements and categorical labels.

Each categorizer label (e.g., fearPB) encodes the viewing angle in its first letter (f, frontal; s, side) and the facial action unit in its final two characters (BL, baseline; PB, ear pulled back; OT, orbital tightening; Bul, nose bulging) (Supplementary Figure 3A). Because the Categorizer classified frontal and lateral views separately, these two views were consolidated into a single measurement for each facial region. Bilateral ear and eye measurements were combined by averaging the two sides when both were available, using the available side when only one was detected, and assigning NaN when neither side was detected. Nose measurements, which comprised a single channel, were used directly.

Two independent analysis streams were then generated.

The behavior-resolved stream (Supplementary Figure 3B), used for Figure 3A-F and Figure 4A, grouped frames according to the Categorizer’s behavioral labels. A frame entered the pain group if either side showed a pain-related facial action unit (PB, OT, or Bul) and the “not pain” group if either side showed baseline (BL). Frames in which the left and right ears or eyes received different labels (“both”) contributed to both groups using the mean of the two measurements. No artifact filtering was applied to this stream. Frames assigned to the “both” category accounted for 4.3% of ear frames and 0.6% of eye frames, whereas the nose measurements were unaffected.

The pain-score stream (Supplementary Figure 3C), used for Figure 4E and Figure 6A-J, followed a different workflow. Individual *intensity_area* measurements were first filtered to remove reflection artifacts and then merged across bilateral facial regions without reference to the Categorizer’s behavioral labels. Consequently, pain-score computation was independent of manual behavioral annotations. The facial action unit-specific weighting factors and Z-score normalization parameters applied at this stage were derived from the CNO dose-response analysis (Figure 4B-D). The resulting merged, filtered measurements served as the input for downstream pain-score computation. When direct comparison between the manual behavior labels and the LabGrymace scores were required, both were displayed simultaneously in the LabGrymace output video (Figure 1N).

#### Reflection artifact filtering

The primary measurement retained for pain scoring was *intensity_area*, together with velocity, magnitude_area, event labels, and XY position. Reflections from the chamber wall occasionally caused the Detector to switch between the true nose position and its mirror image. To remove these artifacts, each tracked channel was filtered before bilateral ear and eye signals were combined, thereby removing transient reflection-driven jumps while preserving genuine facial motion. A frame was classified as a reflection artifact only when all of the following criteria were satisfied: (1) velocity exceeded 10 px/frame; (2) the detected position was within 50 px of the automatically identified mirror location; (3) magnitude_area exceeded 0.5; and (4) at least two additional high-velocity frames occurred within the surrounding 30-frame interval. This procedure removed transient reflection-induced tracking errors while preserving genuine facial movements.

#### Pain score computation and visualization

Filtered ear, eye, and nose intensity_area signals were converted into per-frame normalized values and combined into a composite score using facial region weights derived from the CNO dose-response analysis (W_EAR = 3.574, W_EYE = 3.158, and W_NOSE = 1.689). The composite score was then linearly calibrated to a 0-100 scale, with the uninjected baseline anchored at 0 and the 1 mg/kg CNO condition anchored at 100. For each animal, LabGrymace generated both an overall pain score calculated across all valid frames and windowed pain scores calculated over 3,000-frame windows (100 s at 30 frames/s) to quantify the pain dynamics.

For visualization in the LabGrymace output video, each facial-region intensity was smoothed using a rolling 2-second lookback window. At each frame, the preceding 60 frames were considered, the highest 10% of values were discarded to eliminate tracking spikes, and the remaining values were averaged. The window advanced frame by frame, preserving the temporal resolution of the output. The smoothed per-frame lookback window values were saved to pain_scores_per_frame.xlsx. Because the displayed values are normalized relative to the baseline reference, occasional values below zero simply indicate the estimated pain level fell below the mean baseline value rather than representing negative pain.

### PainFace analysis

#### Model architecture

According to the original PainFace study, PainFace is a cloud-based grimace analysis platform developed for black-coated C57BL/6 mice. Its machine-learning architecture consists of an object-detection component with a ResNet-50 backbone to identify the face/body region, followed by separate convolutional neural networks that assign grimace scores to individual facial action units. For each action unit, the scoring network uses three parallel convolutional branches with kernel sizes of 3 x 3, 5 x 5, and 7 x 7, whose outputs are concatenated and passed to a fully connected classification layer.

#### Input videos and analysis settings

In PainFace, the dark-coated mice setting was selected for all analyses. To match the temporal window used for LabGrymace, only the first 500s of each recording were submitted for analysis. PainFace evaluated videos at 1 frame/s from the original 30 frames/s recordings, corresponding to one analyzed frame every 30 video frames. No additional LabGrymace-style Process Video preprocessing was applied before PainFace analysis, because PainFace was able to directly detect the relevant facial features under its native workflow. However, unlike LabGrymace, PainFace preferentially scores frames in which the face is sufficiently oriented toward the camera for the relevant facial action units to be detected, so frames captured during turning or nonfrontal orientations are less likely to contribute to the final score.

#### Output measures

PainFace outputs were used to obtain both overall grimace scores for each animal and time-resolved grimace scores across the analyzed video interval.

## Statistical analysis

Measurements were exported from LabGym and organized in Excel for manual verification. Subsequent data processing and statistical analyses were performed in Python using pandas and NumPy for data handling, and SciPy for statistical testing, Pearson correlation analysis, and linear regression. Because sample sizes were small, nonparametric tests were used for group comparisons. Wilcoxon signed-rank tests were used for paired comparisons within the same mice. Plots were generated in Python using matplotlib. Data are presented as mean ± SEM. Sample sizes (N) are reported in the corresponding figure legends. Statistical significance is indicated as follows throughout the paper: no significance (ns), p < 0.05 (*), p < 0.01 (**), p < 0.001 (***), and p < 0.0001 (****).

## Data availability

The datasets generated in this study will be deposited to Zenodo and made publicly available.

## Code availability

The code generated in this study is publicly available on GitHub: https://github.com/umyelab/LabGrymace

## Acknowledgements

We thank Robert Tomlinson for helpful discussions, Dr. Bo Duan for sharing reagents, and Julia Philip and Alexa Carr for assisting with the acclimation of mice to the behavioral setup. This research was supported by National Institutes of Health (NIH) grants R01 NS137222 (to B.Y.), UC2 AR082197 and R01 DE035109 (to J.J.E.), and T32 DE007057, T32 DC00011, and K99 DE036046 (to E.A.R.), and University of Michigan Life Sciences Institute Innovation Partnership Fund to B.Y.. The content is solely the responsibility of the authors and does not necessarily represent the official views of funders.

## Author contributions

B.Y., J.E., W.D., K.M.M., and E.A.R conceived the project. B.Y. and J.E. supervised the project. W.D. and B.Y. designed the LabGrymace pipeline and data analysis methods. W.D. programmed the LabGrymace Pain Score module. Y.H. provided inputs on LabGym usage. K.M.M., E.A.R., and J.E. designed and performed the experiments on mice. B.Y., J.E., W.D., and K.M.M. wrote and E.A.R. edited the manuscript.

