## Supplementary Materials for "LabGrymace: Automated Analysis of Mouse Grimace for Quantitative Assessment of Pain Dynamics"

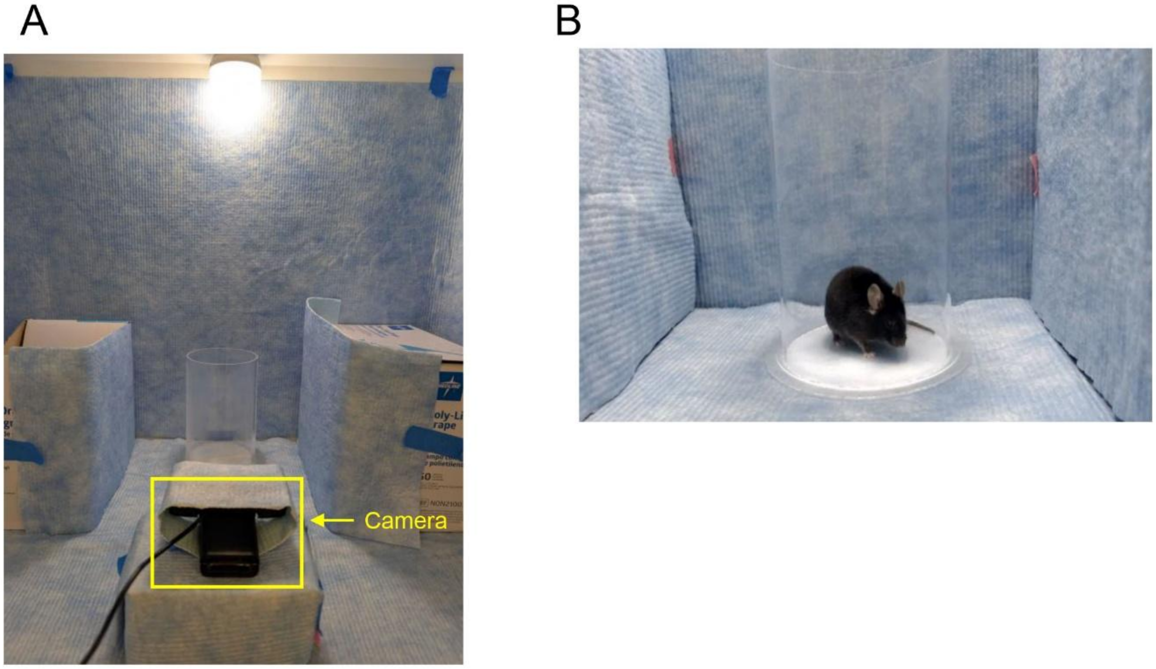


**Supplementary Figure 1. Experimental setup for video recording of mouse facial expressions.**

**(A)** Recording arena. A mouse was placed inside a transparent cylinder enclosure positioned at the center of the arena, allowing visualization of the face while the mouse moved freely. The arena was surrounded by light-blue paper barriers to minimize external visual distractions and provide a uniform background. A camera (yellow box) was positioned at the front of the enclosure to capture the mouse's face, and an overhead light source provided illumination.
**(B)** Representative video frame of a freely moving mouse within the transparent cylinder. Animals were unrestrained and allowed to behave naturally during video recording.
Type or paste caption here. Create a page break and paste in the Figure above the caption.

A

| | precision | recall | f1-score |
|---|---|---|---|
| fEarBL | 0.96 | 0.66 | 0.78 |
| fEarPB | 0.16 | 0.88 | 0.26 |
| fEyeBL | 1.00 | 0.97 | 0.98 |
| fEyeOT | 1.00 | 1.00 | 1.00 |
| noseBL | 0.54 | 0.33 | 0.41 |
| noseBul | 0.72 | 0.85 | 0.78 |
| sEarBL | 0.78 | 0.62 | 0.70 |
| sEarPB | 0.76 | 0.94 | 0.84 |
| sEyeBL | 0.97 | 0.97 | 0.97 |
| sEyeOT | 1.00 | 0.96 | 0.98 |
| accuracy | | | 0.86 |
| macro avg | 0.79 | 0.82 | 0.77 |

B

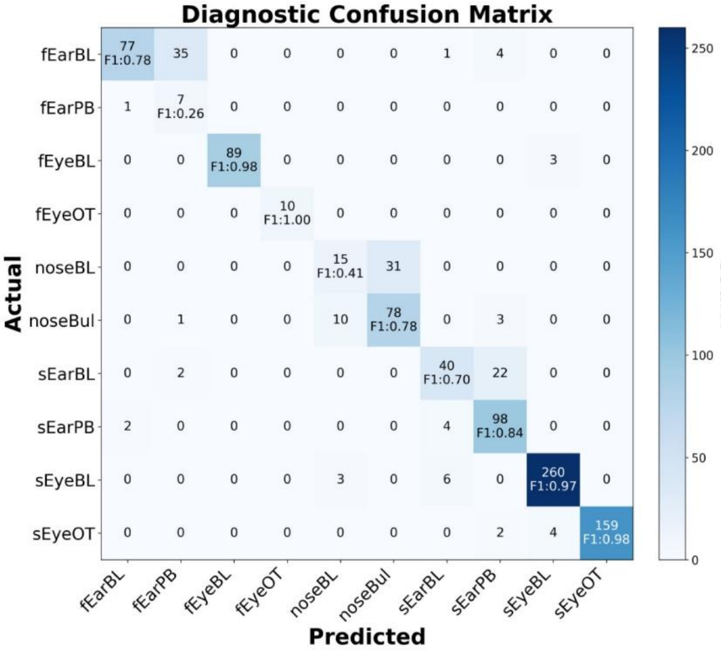


**Supplementary Figure 2. Performance of the facial-action Categorizer on the manually labeled testing set.**

**(A)** Per-class classification performance of the facial-action Categorizer, including precision, recall, and F1 score for each of the ten facial action categories. Performance was evaluated using a manually labeled $MgSO_4$ testing dataset comprising 967 examples.
**(B)** Confusion matrix showing classification performance across the ten facial-action categories. Rows represent manually assigned ground-truth labels, and columns represent predicted labels. Cell color indicates the proportion of testing examples assigned to each predicted category (color scale at right), and diagonal entries display the F1 score for each class. High values along the diagonal indicate accurate classification with minimal confusion between categories.

A

| Time | Ear0View | Ear1View | Ear0Event | Ear1Event | Intensity Area 0 | Intensity Area 1 |
|---|---|---|---|---|---|---|
| 6.46 | lateral | lateral | BL | BL | 5.22 | 5.11 |
| 6.5 | lateral | lateral | BL | BL | 2.42 | 2.82 |
| 6.54 | lateral | lateral | BL | BL | 0.5 | 0.97 |
| 6.58 | lateral | lateral | BL | BL | 1.53 | 0.46 |
| 6.62 | lateral | | PB | | 23.95 | |
| 6.67 | frontal | lateral | PB | PB | 25.62 | 14.93 |
| 6.71 | lateral | lateral | BL | PB | 27.1 | 16.09 |

B

| Time | Ear0Event | Ear1Event | Group | intensity_area |
|---|---|---|---|---|
| 6.46 | BL | BL | not-pain | 5.165 |
| 6.5 | BL | BL | not-pain | 2.62 |
| 6.54 | BL | BL | not-pain | 0.735 |
| 6.58 | BL | BL | not-pain | 0.995 |
| 6.62 | PB | | pain | 23.95 |
| 6.67 | PB | PB | pain | 20.275 |
| 6.71 | BL | PB | both | 21.595 |

C

| Time | IntensityArea_Ear0_filtered | IntensityArea_Ear1_filtered | merged_intensity_area |
|---|---|---|---|
| 6.46 | 5.22 | 5.11 | 5.165 |
| 6.5 | 2.42 | 2.82 | 2.62 |
| 6.54 | 0.5 | 0.97 | 0.735 |
| 6.58 | 1.53 | 0.46 | 0.995 |
| 6.62 | 23.95 | | 23.95 |
| 6.67 | 25.62 | 14.93 | 20.275 |
| 6.71 | | 16.09 | 16.09 |

**Supplementary Figure 3. Combination of facial signals from the LabGym output**

Example illustrating the two analysis streams for the ears of a representative mouse in the CNO dose-response experiment. The left and right ears (Ear0 and Ear1) were tracked independently.

For each frame, the viewing angle (Ear0View/Ear1View: frontal or lateral) and a behavior label assigned by the LabGym categorizer (BL, baseline; PB, pulling back) are shown.
**(A)** Raw frame-level data. Only *intensity_area* is shown for clarity; all nine measurements are represent in the summary sheet and processed identically. Orange marks a reflection artifact.
**(B)** Behavior-resolved stream (used for Figure 3A-F and Figure 4A-D). The two ear measurements are merged for each frame by tanking their mean when both are available, the available value when only one is present, or leaving the value empty when neither is detected. Frames are then grouped according to the behavioral labels as pain, not-pain, or both (one ear classified as PB and the other as BL).
**(C)** Pain-score stream (used for Figure 4E and Figure 6A-J). Only intensity_area is retained. Reflection artifacts are removed independently from each ear (orange), after which the remaining measurements are merged using the same rule as in (B) to generate "merged_intensity_area" (yellow). Frames in which both ear measurements are removed remain empty. The merged values are subsequently used for pain-score computation without reference to the behavioral labels.
